# Physiologically relevant miRNAs in mammalian oocytes are rare and highly abundant

**DOI:** 10.1101/2021.08.08.455568

**Authors:** Shubhangini Kataruka, Veronika Kinterova, Filip Horvat, Jiri Kanka, Petr Svoboda

## Abstract

miRNAs, ~22nt small RNAs associated with Argonaute (AGO) proteins, are important negative regulators of gene expression in mammalian cells. However, mammalian maternal miRNAs show negligible repressive activity and the miRNA pathway is dispensable for oocytes and maternal-to-zygotic transition. The stoichiometric hypothesis proposed that this is caused by dilution of maternal miRNAs during oocyte growth. As the dilution affects miRNAs but not mRNAs, it creates unfavorable miRNA:mRNA stoichiometry for efficient repression of cognate mRNAs. Here we report that porcine *ssc-miR-205* and bovine *bta-miR-10b* are exceptional miRNAs, which resist the diluting effect of oocyte growth and can efficiently suppress gene expression. Additional analysis of *ssc-miR-205* shows that it has higher stability, reduces expression of endogenous targets, and contributes to porcine oocyte-to-embryo transition. Consistent with the stoichiometric hypothesis, our results show that the endogenous miRNA pathway in mammalian oocytes is intact and that maternal miRNAs can efficiently suppress gene expression when a favorable miRNA:mRNA stoichiometry is established.

## Introduction

MicroRNAs (miRNAs, reviewed in detail in [1]) are genome-encoded ~22 nt long single-strand small RNAs, which guide post-transcriptional repression of gene expression. A typical mammalian miRNA biogenesis (reviewed in [2]) involves nuclear processing of primary miRNA transcripts (pri-miRNA) into small hairpin precursors (pre-miRNA), which are transported to the cytoplasm. A cytoplasmic pre-miRNA is cleaved by RNase III Dicer and a miRNA is loaded onto an Argonaute protein, the key protein component of the effector complex named RISC (RNA-Induced Silencing Complex). Mammalian genomes encode four AGO proteins, which accommodate miRNAs equally well [3] and appear to be functionally redundant in the miRNA pathway [4]. However, AGO2 stands out among the mammalian AGO proteins, as it carries endonucleolytic activity cleaving cognate RNAs perfectly complementary to a loaded small RNA [3,5]. AGO2 with a small RNA thus forms the minimal RISC (holo-RISC), which traditionally has been associated with the RNA interference pathway [6]. While miRNAs guide RNAi-like endonucleolytic cleavage of perfectly complementary mRNAs by AGO2 as well [7,8], a typical miRNA:mRNA interaction occurs through imperfect complementarity involving the “seed” region comprising nucleotides 2 to 8 of the miRNA [9,10]. Target repression through this type of interaction, referred to miRNA-like hereafter, is slower as it involves weaker and longer association of RISC with the target [11,12]. It also involves additional proteins, which form the full miRNA-loaded RISC (miRISC). The key AGO2-binding partner is GW182 adaptor protein, which recruits further protein factors mediating translational repression coupled with deadenylation and decapping [13–16].

miRNAs were implicated in countless physiological processes and pathologies. Thousands of mammalian miRNAs were annotated [17] and more than a half of mammalian genes could be directly targeted by miRNAs [18]. Yet, miRNAs are dispensable for mouse oocytes and preimplantation development and their activity in oocytes is negligible [19,20]. Notably, analysis of miRNAs in mammalian oocytes revealed their low cytoplasmic concentration (<0.5 nM) [21] as opposed to functional miRNAs in somatic cells, which are relatively abundant [22–24]. This is consistent with kinetic studies highlighting miRNA concentration as an important factor for efficient miRNA-mediated repression [11,12]. For example, analysis of *let-7* miRNA in HeLa cells, whose transcriptome size was estimated to be ~580,000 mRNAs molecules/cell [25], determined that a HeLa cell contains ~50,000 *let-7* molecules [22]; this equals to ~20 nM cytoplasmic concentration (assuming ~20 μm cell diameter). In contrast, the cytoplasmic mRNA concentration in somatic cells and oocytes is comparable despite somatic cells containing 0.1-0.6×10^6^ mRNA molecules while mouse oocytes accumulate a sizeable transcriptome of ~27×10^6^ mRNA molecules [25–30]. Maintenance of mRNA concentration in the cytoplasm in oocytes correlates with extended average half-life of maternal mRNAs [31–33] while turnover of maternal miRNAs does not appear to be adapted to the oocyte growth resulting in their dilution [21].

The “stoichiometric model” explaining the loss of physiologically significant activity of the miRNA pathway thus proposes that oocyte’s growth dilutes maternal miRNA concentration to the point where miRNA become ineffective regulators of the maternal transcriptome. Another explanation was offered by Freimer et al. who proposed that alternative splicing of *Ago2* makes the functional AGO2 a limiting factor, which contributes to the observed miRNA inactivity [34]. However, although this mouse-specific *Ago2* regulation could contribute to the negligible miRNA activity in mouse oocytes, it cannot explain low miRNA abundance and inactivity observed in bovine and porcine oocytes [21]. Here we report identification and analysis of two exceptionally abundant maternal miRNAs in bovine and porcine oocytes, which overcome the diluting effect, exhibit robust repressive activity, and support the stoichiometric model.

## Results and Discussion

### Extreme abundance of ssc-miR-205 and bta-miR-10b in oocytes

During analysis of published small RNA sequencing (RNA-seq) data [35,36], we noticed exceptional abundance and possible functional relevance of *ssc-miR-205* and *bta-miR-10b* in porcine and bovine oocytes, respectively (Fig. 1A). Even though *miR-205* and *mir10b* are conserved across vertebrates [37], their high maternal expression is not conserved in mammals (Fig. 1A). There are minimal sequence differences among the mouse, porcine, and bovine *miR-205* and *mir10b* miRNA precursors (Fig. 1B). Consequently, the secondary structure of *ssc*-*miR-205* and *bta*-*miR-10b* seems preserved and thus not associated with their high abundance in porcine and bovine oocytes, respectively (Fig. 1B and S1A).

**Figure 1.**
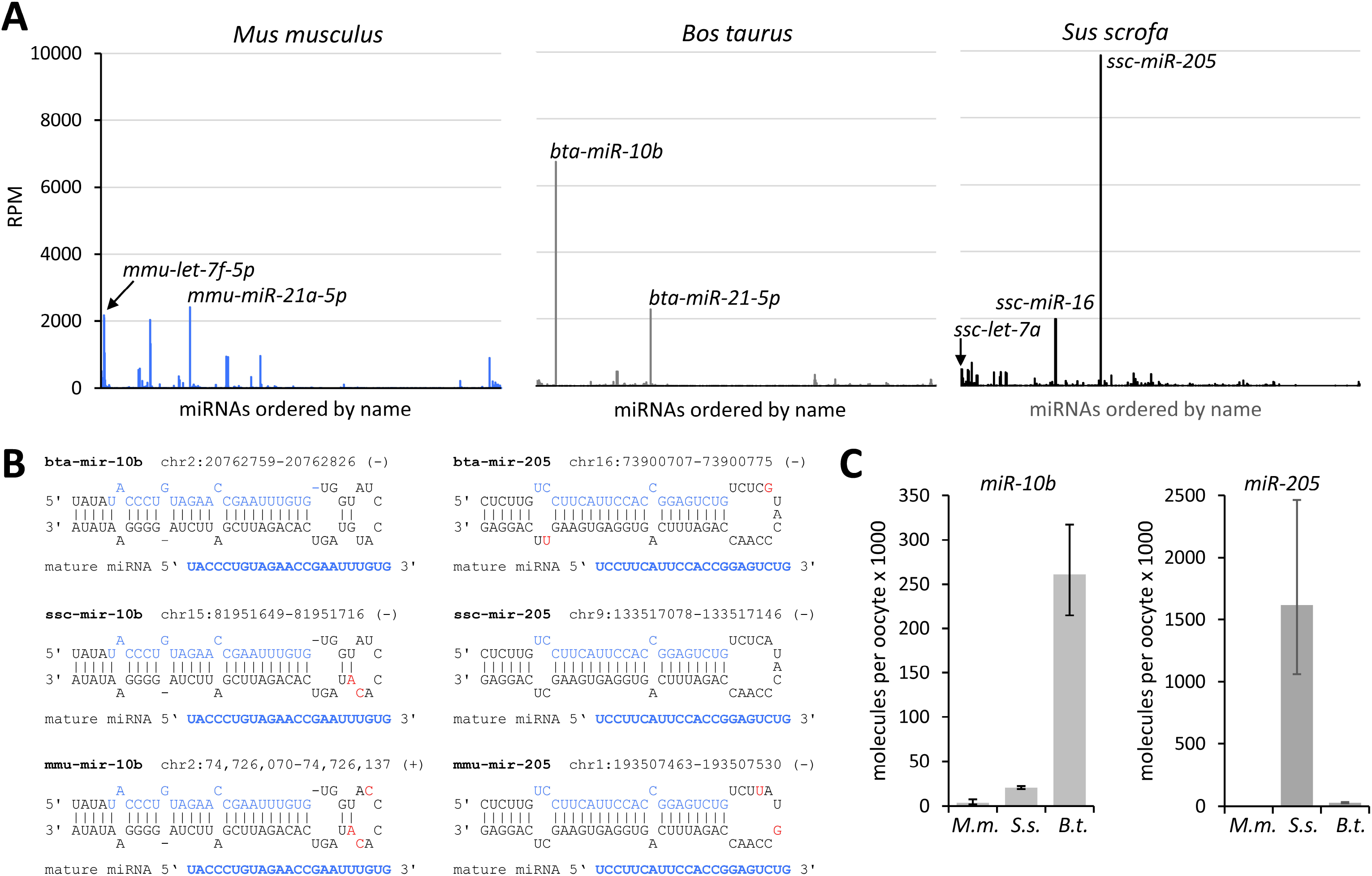
Functional miRNAs in mammalian oocytes (A) miRNA abundance in RNA-sequencing samples from murine, porcine, and bovine oocytes [35,53,54]. Each graph shows on the y-axis reads per million of 19-32 nt reads for miRNAs and on the x-axis miRNAs ordered by name. (B) Schematic depiction of miRNA precursors. Precursor sequences and predicted basepairing was taken from miRbase miRNA annotations [17]. *miR-10b* secondary structures in the mirRbase contained two different basepairing versions in the loop, the alternative folding is displayed in Fig. S1A. (C) qPCR quantification of *ssc-miR-205* and *bta-miR-10b* miRNAs in murine (*M.m.*), porcine (*S.s.*), and bovine (*B.t.*) oocytes, respectively.

Since RNA-seq data provided only relative estimates of miRNA abundance, we used quantitative RT-PCR to determine copy numbers per oocyte (Fig. 1C): *ssc-miR-205* was estimated to have ~1,6 million molecules per oocyte (~4.4 nM cytoplasmic concentration) and *bta-miR-10b* ~261,000 molecules per oocyte (~0.5 nM cytoplasmic concentration). Concentrations were estimated using 105 μm and 120 μm diameters for porcine and bovine oocytes, respectively [38]. In mouse oocytes, we have observed functional repression by miRNAs at 1.5 nM but not at 0.3 nM concentration [21]. Thus endogenous *ssc-miR-205* would be predicted to suppress gene expression in oocytes while repressive potential of *bta-miR-10b* was unclear.

### Endogenous ssc-miR-205 and bta-miR-10b are active in oocytes

To estimate *ssc-miR-205* and *bta-miR-10b* activities in oocytes, we produced and microinjected luciferase reporters with perfectly complementary (“perfect”), partially complementary (“bulged”), or mutated miRNA binding sites for *miR-205* or *miR-10b* (Fig. S1B). Perfect and bulged reporters allow for partially distinguishing between RNAi-like cleavage of the target and typical miRNA-mediated repression [8]. Importantly, for miRNA-targeted reporters, we used NanoLuc luciferase [39], arguably the best tool for determining endogenous miRNA activity in oocytes. NanoLuc allows usage of approximately 10,000 reporter mRNA molecules per oocyte [21], which is well within the physiological range of maternal mRNAs [30].

Both miRNAs efficiently repressed perfect and bulged reporters demonstrating that *ssc-miR-205* and *bta-miR-10b* are active and efficiently repressing targets (Fig. 2A). Consistent with its much higher abundance, *ssc-miR-205* in porcine oocytes attained much stronger reporter repression than *bta-miR-10b* in bovine oocytes (Fig. 2A). These results are the strongest evidence for efficient target repression by endogenous miRNAs in mammalian oocytes so far.

**Figure 2.**
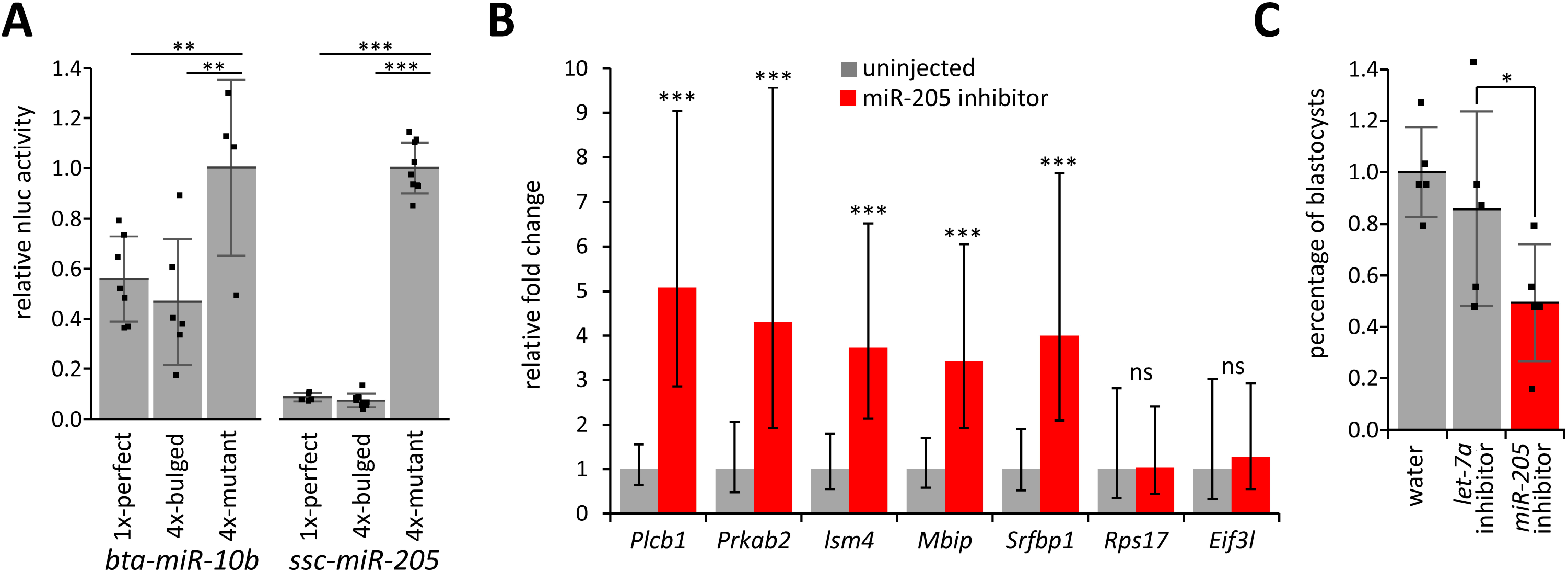
(A) *bta-miR-10b* and *ssc-miR-205* miRNAs efficiently repress microinjected NanoLuc luciferase reporters carrying a perfectly complementary (1x perfect) miRNA binding site or four partially complementary (bulged) miRNA binding sites. See the supplementary material for more information about the reporter design. (B) De-repression of predicted *ssc-miR-205* targets. *ssc-miR-205* was inhibited by a microinjected antisense oligonucleotide inhibitor and target levels were assessed 24 hours after microinjection by RT-qPCR. Error bars = SD. Asterisks indicate statistical significance (p-value) of one-tailed *t*-test, (* < 0.05, ** < 0.01, *** < 0.001) for (A) and (B) (C) Inhibition of *ssc-miR-205* by a microinjected antisense oligonucleotide inhibitor results in reduced development to the blastocyst stage. Shown is development of microinjected oocytes to the blastocyst stage relative to water-injected oocytes. Five independent experiments (represented by individual data points) were performed, ~50 porcine oocytes were microinjected in each group in each experiment. Difference between *let-7a* and *miR-205* inhibitors effects is statistically significant (two-tailed paired t-test p-value = 0.016). Error bars = SD.

miRNA activities in mammalian oocytes do not appear to be suppressed in *sensu stricto*. The miRNA pathway appears mechanistically intact but cytoplasmic miRNA concentrations are much lower than in somatic cells and consequently, miRNA targets are repressed weakly. It is important that analysis of miRNAs in oocytes and zygotes would address and respect physiological concentrations of miRNAs and their targets as experimental conditions beyond physiological ranges would increase the risk of producing non-physiological artifacts.

Several studies reported presence of functional miRNAs in murine, bovine or porcine oocytes [40–43]. In these studies, experimental support for significant endogenous miRNA activity in oocytes was inferred from correlative effects of miRNA overexpression or use of antisense inhibitory oligonucleotides (“antagomirs”) [44]. For example, Chen et al reported that *miR-27a* activity is not suppressed in porcine oocytes [40]. However, their functional analysis employed microinjection of oocytes with 50 pl of 500 ng/μl of miRNA mimics or inhibitors. That would correspond to more than a billion microinjected oligonucleotides while an oocyte contains tens of millions of mRNA molecules [30]. Similarly, *miR-98* function in mouse oocytes was studied using microinjection of 10 pl of 50 μM miRNA or 100 μM inhibitors [41]. Since cytoplasmic volume of a mouse oocyte is ~260 pl, such experimental design would result in micromolar concentrations of microinjected oligonucleotides in the cytoplasm. This is three orders of magnitude or more than estimated amounts of maternal miRNAs [21] and an order of magnitude excess over the amount of maternal mRNAs [30]. The analysis of *miR-130b* in bovine oocytes was conducted near the physiological range, as 10 pl of 50 nM miRNA or inhibitors were injected [42]. At the same time, this injected amount is higher than the amount of *bta-miR-10b*, the most abundant miRNA in bovine oocytes reported here and there is no evidence that bta-*miR-130b* would reach similar abundance. In fact, miR-130b has three orders of magnitude lower abundance than miR-10b in RNA-seq from bovine oocytes [36].

### Endogenous ssc-miR-205 is biologically relevant

After determining the concentrations of *ssc-miR-205* and *bta-miR-10b* and assessing their activity, we focused on subsequent functional analysis of *ssc-miR-205*, which showed stronger target repression and porcine oocytes were easily accessible for experimental analysis. To test whether *ssc-miR-205* suppresses endogenous targets, we inhibited the miRNA in the oocyte with *miR-205* antagomir microinjection and examine the mRNA levels of five predicted endogenous targets by qPCR. The selected targets *Plcb1*, *Prkab2*, *Lsm4*, *Mbip* and *Srfbp1* were the best predicted *ssc-miR-205* targets in miRBase [17] and their expression was detected in RNA-seq data from porcine oocytes [35].

To test suppression of the endogenous targets, we microinjected fully-grown transcriptionally quiescent porcine oocytes [45] with ~5×10^6^ molecules of *miR-205* antagomir. This amount corresponds to ~2-fold excess over endogenous *ssc-miR-205* molecules and is much lower than amounts used to study miRNAs in oocytes in the studies mentioned above [30,40]. Microinjected fully-grown oocytes were cultured for 24 hours in the presence of dbcAMP, which prevents resumption of meiosis [46]. As the first wave of maternal mRNA degradation is induced during resumption of meiosis [47], the use of dbcAMP allows to avoid interference of complex meiotic mRNA degradation with monitoring target degradation by miRNAs.

*miR-205* antagomir caused a significant 3-5 fold increase in mRNA abundance for all predicted targets but not for two non-targeted control genes (Fig. 2B). De-repression of the endogenous targets upon injection of *miR-205* antagomirs complements results from NanoLuc reporter experiments and implies that *ssc-miR-205* is indeed functional in porcine oocytes and regulating endogenous gene expression in porcine oocytes.

Next, we examined whether *ssc-miR-205* plays a significant biological role in oocytes and/or early embryos. We microinjected fully-grown porcine oocytes with the *miR-205* antagomir (~2-fold excess over endogenous *ssc-miR-205* molecules) and allowed them to undergo meiotic maturation, and then parthenogenetically activated their preimplantation development. We opted for parthenogenetic early development because porcine oocytes suffer a high incidence of polyspermy [48] while parthenogenotes can progress through the preimplantation development to the blastocyst stage [49]. As a negative control, we used *let-7a* inhibitor as this miRNA is present but not effective in porcine oocytes [21].

*ssc-miR-205* inhibitor-injected oocytes showed ~50% reduced ability to support development to the blastocyst stage relative to water-injected control and *let-7a* inhibitor-injected oocytes (Fig. 2C). The precise role of miR-205 inhibition in the phenotype remains unknown. There was no specific stage at which the development of *miR-205* antagomir injected oocytes arrested and there are many predicted *ssc-miR-205* targets. The miRNA could play a role in shaping the zygotic expression or contribute to maternal mRNA degradation. Limitations of the model prevent determining whether *miR-205* antagomir injected blastocysts would develop normally or show additional defects, similarly to *miR-430* in zebrafish. *miR-430* is uniquely adapted for rapid and massive expression upon fertilization and contributes to maternal mRNA degradation in zebrafish embryos [50]. Deletion of the *miR-430* cluster in zebrafish causes developmental delay and morphological defects at earlier developmental stages while mutant embryos are able to undergo gastrulation and organogenesis before they die five days after fertilization [51]. Thus, dissection of the biological role of *ssc-miR-205* in early porcine embryos will require further analysis of *ssc-miR-205* effects on maternal and zygotic transcriptomes to reveal the phenotype severity and strengthen the link between the phenotype and *ssc-miR-205* function.

### High abundance of ssc-miR-205 correlates with its slower turnover

Exceptional abundance of *ssc-miR-205* in porcine oocytes suggests a unique adaptation, which enables its accumulation during porcine oocyte growth but does not exist in bovine and murine oocytes. As mentioned above, there are minimal differences among the porcine, bovine, and murine miRNA precursors which do not have any apparent effect on the secondary structure of the precursor (Fig. 1B). This makes unique secondary structure or sequence variability of mature miRNA or miRNA* less likely explanations for *ssc-miR-205* accumulation.

Porcine oocyte-specific high abundance of *ssc-miR-205* could come from unusually high transcription of the miRNA precursor. However, RNA-sequencing from mammalian oocytes [35,36,52–55] does not support unusually high transcription at the *ssc-miR-205* locus (Fig. 3A).

**Figure 3.**
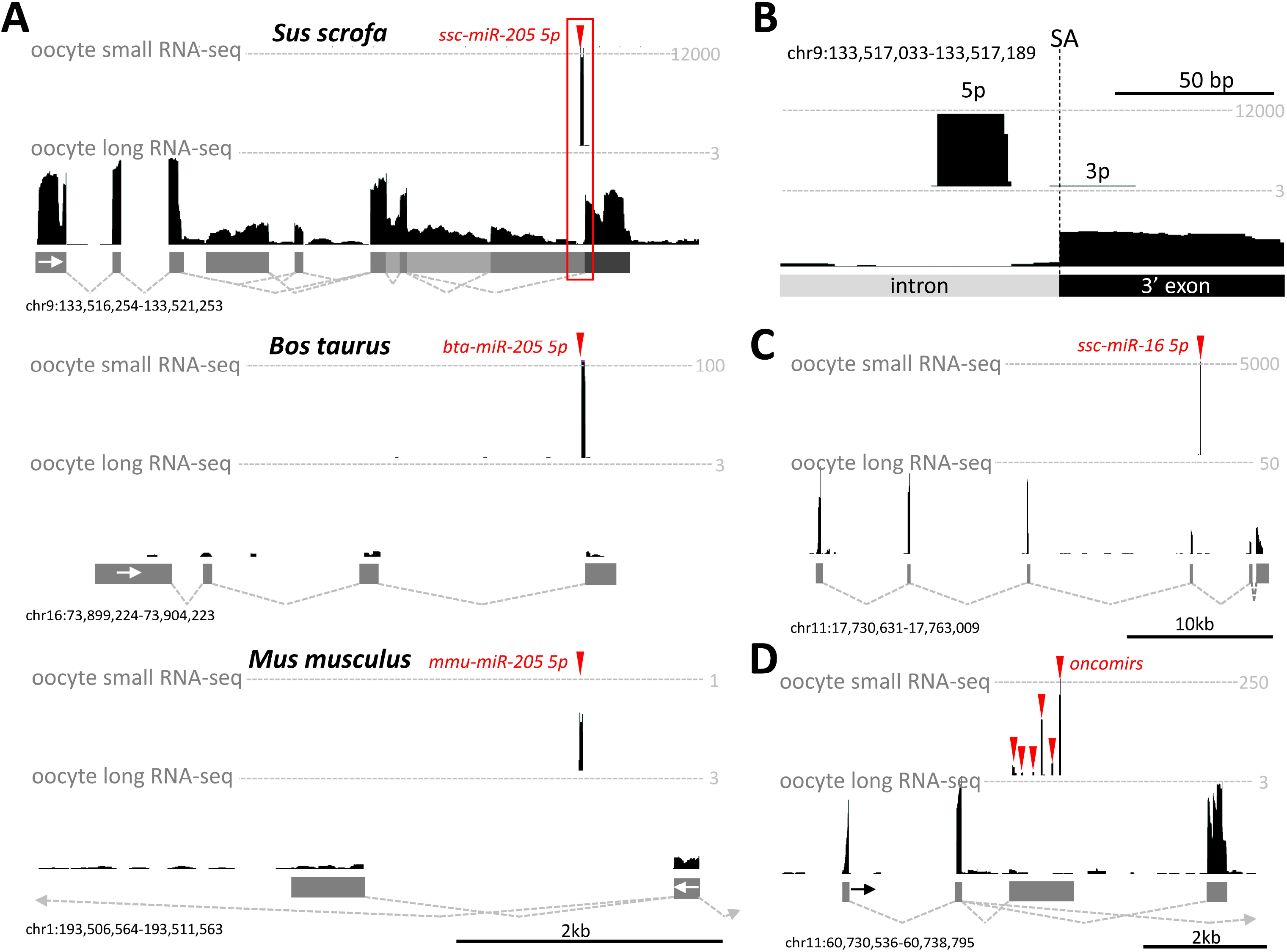
UCSC browser snapshots depicting transcription in selected miRNA loci from selected species. Tracks from small and long RNA-seq analyses of mouse [54,55], bovine [36,53], and porcine oocytes [35,52] were constructed as described in material and methods. The y-scale for small RNAs depicts counts per million (CPM) of mapped 19-32 nt reads. The y-scale for long RNA-seq depicts counts per million of mapped fragments. Grey rectangles and dashed lines represent exon-intron structure and splicing of transcripts in the locus inferred from RNA-seq data. All UCSC browser snapshots are oriented such that depicted miRNAs are transcribed from the left to the right. (A) A UCSC browser snapshot of the *miR-205* locus depicting small and long RNA expression in selected loci in porcine, bovine, and mouse oocytes. Despite the splice acceptor of the terminal exon appears conserved, RNA-seq data do not support splicing of the primary *miR-205* transcript. Instead, the murine miR-205 locus produces a detectable antisense spliced maternal lncRNA while the precursor of mmu-miR-205 is undetectable. (B) A detail of the *ssc-miR-205* locus marked in the panel A by the red rectangle. SA, splice acceptor for the last exon of lncRNA originating from the locus. (C) A UCSC browser snapshot of the *ssc-miR-16 locus* giving rise to the second most abundant miRNA in the sequencing dataset [35]. (D) A UCSC browser snapshot of the porcine *miR-17/92* (oncomir cluster) locus.

Porcine and bovine *miR-205* apparently originate from a nascent transcript of a spliced lncRNA (Fig. 3A). Notably, production of the pre-miRNA and splicing of the lncRNA are mutually exclusive because of intron/exon boundary within the pre-miRNA (Fig. 3B). Albeit the boundary appears conserved in mice, RNA-seq data from oocytes and somatic tissues do not provide evidence for splicing of the murine *miR-205* locus [55, Abe, 2015 #124,56,57] (Fig. 3A). Expression of the lncRNA is well detectable in porcine oocytes when compared to bovine oocytes, which show much lower expression of both, *bta*-*miR-205* and the lncRNA (Fig. 3A). However, RNA amounts observed in RNA-seq data in the porcine locus are rather low (<3CPM) to explain production of 1.6 million *ssc-miR-205* molecules per oocyte. In comparison, highly abundant mRNAs such as *Zp3* and *Gdf9*, which would be expected to have hundreds of thousands of mRNA copies/oocyte, have expression around 700-800 CPM. Also, comparison with other porcine miRNA loci shows similar or even higher expression of spliced lncRNA precursor. For example, the *ssc-miR-16* locus produces the second most abundant maternal miRNA. The miRNA is localized in an intron of a spliced lncRNA, which has much higher abundance than the spliced lncRNA at the *ssc-miR-205* locus (Fig. 3C). A similar abundance to the lncRNA from the *ssc-miR-205* locus was found for a spliced lncRNA transcript from the *miR-17-92* cluster locus (also known as the “OncomiR” cluster [58]). However, relative to *ssc-miR-205,* the six miRNAs from the oncomir cluster had lower amounts with considerable variability among the oncomirs (Fig. 3D). Taken together, RNA-seq data from the porcine *miR-205* locus suggest that sole transcription of the *ssc-miR-205* pri-miRNA is not the primary underlying cause of the high accumulation of the mature miRNA.

We also investigated whether *ssc-miR-205* accumulation could be regulated at the level of the mature miRNA. We first examined decay of *ssc-miR-205* in fully grown oocyte since miRNA turnover appeared to be a contributing factor to miRNA dilution in mouse oocytes. We cultured fully-grown oocytes in the presence of dbcAMP for 10, 20, and 40 hours and observed that *ssc-miR-205* exhibits higher stability than *ssc-let-7a* (Fig. 3E). *Let-7a* belongs to medium-to-high abundant miRNAs in porcine oocytes (Fig. 1A) [35]. While *ssc-let-7a* levels were reduced by ~70% over the course of 40 hours (similarly to mouse oocytes [21]), *ssc-miR-205* levels were reduced only by ~25% after 40 hours (Fig. 4A). This suggests that accumulation of *ssc-miR-205* during oocyte growth involves increased miRNA stability.

**Figure 4.**
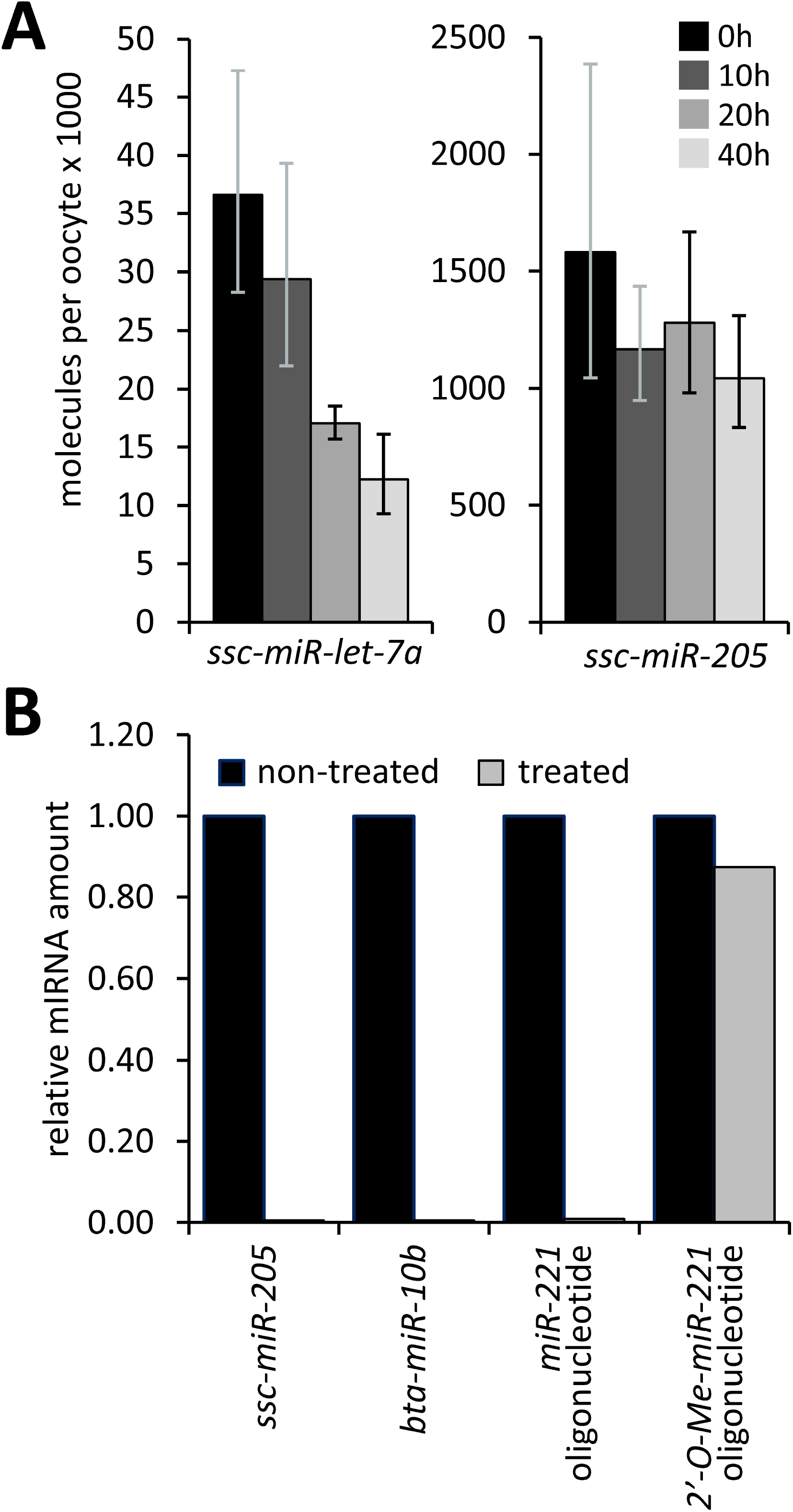
(A) *ssc-miR-205* has increased stability in porcine oocytes. Fully-grown transcriptionally quiescent oocytes were cultured for indicated periods of time and the amount of *ssc-let-7a* and *ssc-miR-205* was estimated by RT-qPCR. All error bars represent standard deviation (SD). (B) Oxidation by sodium periodate followed by qPCR does not support 2’-OH modification of either ssc-miR-205 or bta-miR-10b miRNA. Shown are relative levels of miRNAs with and without the sodium periodate treatment. Non-methylated and methylated miR-221 RNA oligonucleotides were used as a positive control for the assay.

The cause of increased stability of *ssc-miR-205* remains unclear but it does not involve a 2’-OH modification of *ssc-miR-205* (Fig. 4B), such as 2’-O-methylation, which is common for plant miRNAs and was reported to stabilize *miR-21-5p* in lung cancer [59]. Unlike the positive control (2’-methylated *miR-221*), which remained well detected upon the periodate treatment (Fig. 4B), endogenous *ssc-miR-205* and *bta-miR-10b* miRNAs vanished upon the treatment suggesting that they do not carry 2’-O-methylation.

A good candidate mechanism for *ssc-miR-205* accumulation is interaction with some maternal RNA molecule, which would increase *ssc-miR-205* stability. Such an RNA could explain species-specific and tissue-specific *ssc-miR-205* accumulation in porcine oocytes. A precedent for such a regulation could be effect of circular RNA *Cdr1* on *miR-7* in the mouse brain [60]. However, search for a putative binding partner among circRNAs, lncRNA, and mRNAs did not yield any outstanding candidate, which would have high abundance and multiple *ssc-miR-205* binding sites. The strongest candidate was a highly abundant unspliced ~28 kb long piRNA precursor RNA carrying several *miR-205* seed motifs. However, given the length of this RNA, it is hard to envision how this transcript could selectively stabilize *ssc-miR-205* and not other maternal miRNAs.

Taken together, we show that highly abundant maternal miRNAs *ssc-miR-205* and *bta-miR-10b* overcome the constraints imposed on miRNA activity by the size of the maternal transcriptome and volume of the oocyte. Our results support the model that the main cause of the apparent maternal miRNA inactivity is unfavorable miRNA:mRNA stoichiometry. In addition, we provide a framework for the identification of functionally relevant miRNAs in the oocyte. *ssc-miR-205* reported here is the best experimentally documented example of an active maternal mammalian miRNA, which functionally contributes to zygotic development. At the same time, the lack of conserved high abundance of *miR-205* and *miR-10b* in mammalian oocytes suggests that these two miRNAs represent unique evolutionary events. Thus, they are exceptions from the common biological insignificance of maternal miRNAs that did not adapt to the diluting effect of oocyte growth.

## Material and Methods

### Oocyte collection and microinjection

Porcine and bovine oocytes were obtained from the slaughterhouse material as described previously [61,62]. For reporter injection, a mixture of *in-vitro* transcribed firefly and nanoluciferase (NanoLuc) RNA in the ratio of 100,000:10,000 was injected per oocyte with FemtoJet microinjector (Eppendorf). For antisense oligonucleotide inhibitor injection, commercially obtained hsa-let-7a-5p or hsa-miR-205-5p miRCURY LNA miRNA Inhibitor (Qiagen, cat# YI04101776 and YI04101508, respectively) was diluted in water and microinjected (~5×10^6^ molecules per oocytes).

Injected bovine oocytes were cultured in MPM media (prepared in house [62]) containing 1 mM dbcAMP (Sigma) without a paraffin overlay in a humidified atmosphere at 39 °C with 5% CO_2_ for 20 h. Porcine oocytes were cultured in M-199 MEDIUM (Gibco) supplemented with 1 mM dbcAMP, 0.91 mM sodium pyruvate, 0.57 mM cysteine, 5.5 mM Hepes, antibiotics and 5% fetal calf serum (Sigma). Injected oocytes were incubated at 38.5 °C in a humidified atmosphere of 5% CO_2_ for 20 h.

### Parthenogenetic activation and embryo culture

After maturation, oocytes were washed twice in PXM-HEPES and activated by exposure to 10μM ionomycin in PXM-HEPES for 5 min. This was followed by two washes in porcine zygote medium 3 (PZM 3) supplemented with 2 mM 6-DMAP and cultivation for 5 h at 38.5°C in 5% CO2. Then they were washed twice in PZM 3 and cultivated for 7 more days in PZM 3 medium.

### cDNA synthesis and qPCR

The oocytes were washed with M2 media to remove any residual IBMX, collected in minimum amount of M2 media with 1 μl of Ribolock, and incubated at 85 °C for 5 minutes to release the RNA. cDNA synthesis for miRNA analysis was done with the miRCURY LNA RT kit (Qiagen, cat# 339340) according to the manufacturer’s protocol. qPCR was set using cDNA fraction corresponding to one oocyte equivalent per qPCR reaction. The miRCURY LNA SYBR green kit (Qiagen. cat# 339345) was used as per manufacturer’s protocol and the reaction was set in Roche LightCycler 480. The following primer sets from Qiagen were used: hsa-let-7a-5p (cat# YP00205727), hsa-miR-205-5p (cat# YP00204487), hsa-mir-10b-5p (cat# YP00205637), hsa-mir-221-3p (cat# YP00204532).

For mRNA analysis, oocytes were washed and lysed for cDNA synthesis as for miRNA. Reverse transcription with Maxima Reverse transcriptase (Thermo scientific) was primed with random hexanucleotides. Maxima SYBR Green qPCR Master Mix (Thermo Scientific) was used for qPCR. qPCR was set using cDNA fraction corresponding to one oocyte equivalent per qPCR reaction in technical triplicates for each biological sample (also in triplicates). Average CT values of the technical replicates were normalized to uninjected control oocytes.

### Plasmid reporters

Nanoluciferase plasmid pNL1.1 was obtained from Promega. The Nluc gene was cleaved out and ligated in the phRL-SV40 plasmid downstream of the T7 promoter. miRNA binding sites were obtained as oligonucleotides from Sigma. Oligonucleotides were annealed, phosphorylated and cloned downstream of the Nluc gene in XbaI site. The firefly control plasmid was made in house with a T7 promoter (oligonucleotide sequences are provided in the Table S1).

### In vitro transcription

Linearized pNL-miR-205 and miR-10b-perfect, bulged, mutant and firefly plasmids were *in vitro* transcribed, capped and then polyadenylated by Poly(A) tailing kit (ThermoFisher).

### Luciferase assay

Five oocytes were collected per aliquot and Nano-Glo Dual-Luciferase reporter assay system (Promega, cat# N1610) was used to measure the samples as per the manufacturer’s protocol.

### 2’-OH methylation assay

Oligonucleotides or oocytes (supplemented with a methylated miR-221 oligonucleotide as an internal normalization control) were incubated in 50 μl Borax buffer with sodium periodate for 30 min at room temperature in dark. Next, 3 μl of 60% glycerol was added and further incubated for 10 min at RT. This was followed by phenol-chloroform extraction and precipitation with isopropanol. The pellet was dissolved in water and used for cDNA synthesis followed by qPCR, which was set in technical triplicates for each biological sample (also in triplicates). Average CT values were first normalized to internal methylated miR-221 control followed by normalization to untreated sample.

### Analysis of high throughput sequencing data

Small RNA-sequencing data from mouse (GSE59254 [54]), porcine (GSE122032 [35]), and bovine (GSE 64942 [36]) oocytes were mapped to their respective genomes (mouse – mm10, cow – bosTau9, pig – susScr11) as described previously [63]. After mapping, 21-23 nt long reads perfectly aligned to the genome were selected for mature miRNA expression quantification with annotation from miRBase [17] using featureCounts v2.0.0 program [64]:

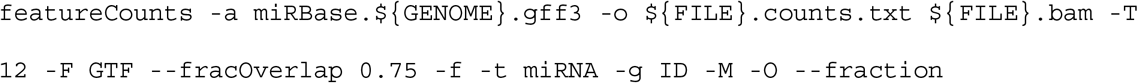

For bovine and porcine samples, miRBase annotation was lifted-over from older genome versions using Liftoff [65].

Long RNA-seq data from mouse (GSE116771 [55]), porcine (PRJNA548212 [52]), and bovine (GSE52415 [53]) fully-grown oocytes and were mapped to their respective genomes (mouse – mm10, cow – bosTau9, pig – susScr11) using STAR 2.5.3a [66] as previously described [55].

Read mapping coverage was visualized in the UCSC Genome Browser by constructing bigWig tracks from merged replicates using the UCSC tools [67].

## Acknowledgements

We thank Radek Malik for help with preparation of the manuscript. This work was funded from the European Research Council under the European Union’s Horizon 2020 research and innovation programme (grant agreement No 647403, D-FENS). IMG institutional support included RVO: 68378050-KAV-NPUI. J.K. and V.K. were in part supported by IAPG institutional support RVO: 67985904. Computational resources for F.H. were supported by European Structural and Investment Funds grant (#KK.01.1.1.01.0010), Croatian National Centre of Research Excellence for Data Science and Advanced Cooperative Systems (#KK.01.1.1.01.0009). Financial support of S.K. and F. H. was in part provided by Charles University in Prague in the form of a PhD student fellowship; this work will be in part used to fulfil the requirements for a PhD degree and hence can be considered “school work”.

## Author contributions

S.K. and V.K. performed the experiments. F.H. analyzed RNA-sequencing data. S.K, J.K., and P.S. designed the experiments, supervised, and analyzed experimental data. S.K. and P.S. wrote the manuscript.

---

**Figure S1** Additional schemes of pre-miRNAs and reporter designs (A) Alternative fold of bovine, porcine, and murine *miR-10b*. The fold was predicted in miRbase for bta-mir-10b and the remaining miRNAs were folded accordingly. (B) Schematic depiction of NanoLuc reporters and of sequences of miRNA binding sites.

**Supplementary Table 1.**
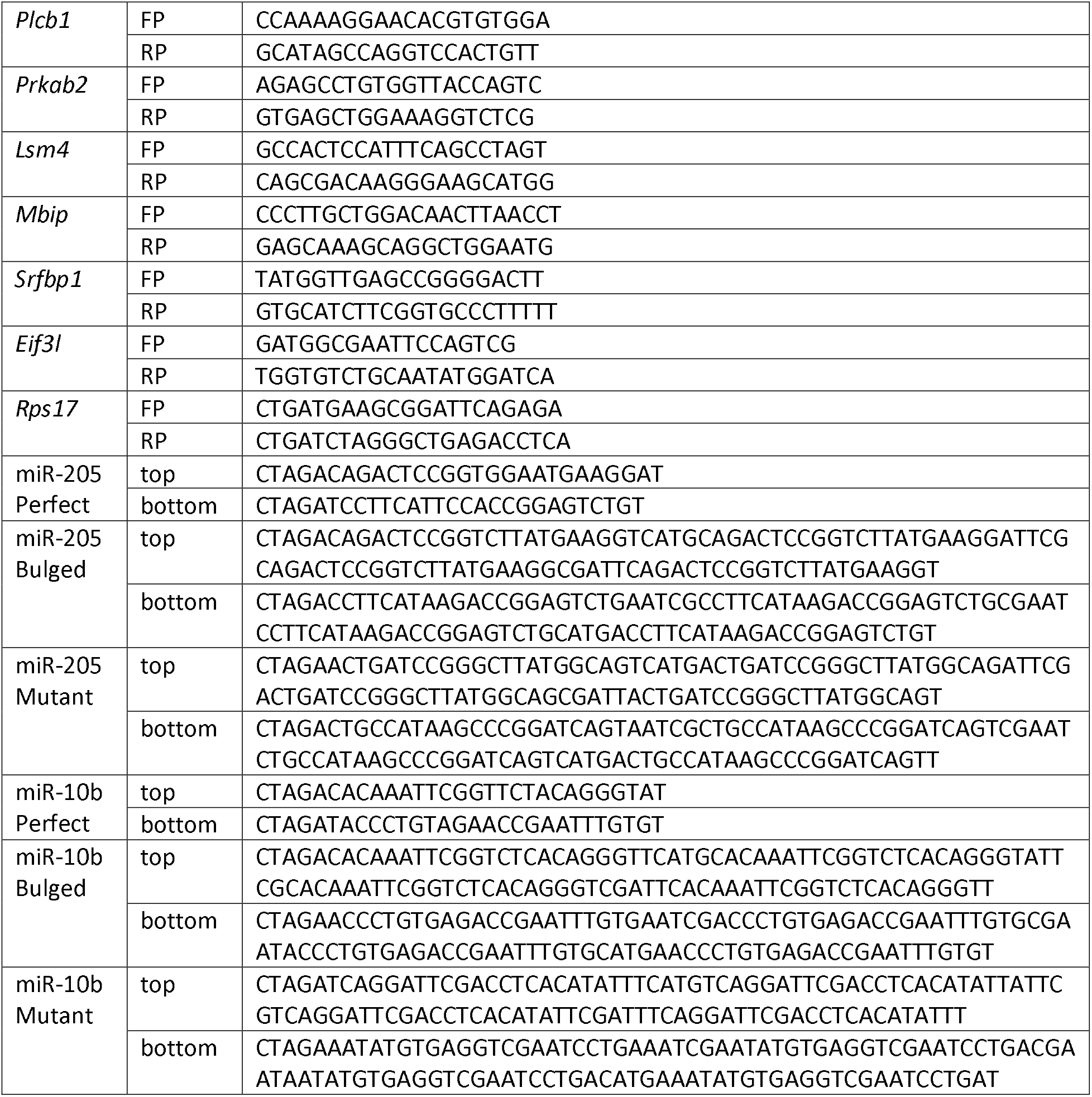
Primers and oligonucleotides

