## Supplementary figures and images for "Physiologically relevant miRNAs in mammalian oocytes are rare and highly abundant"

### Figure S1

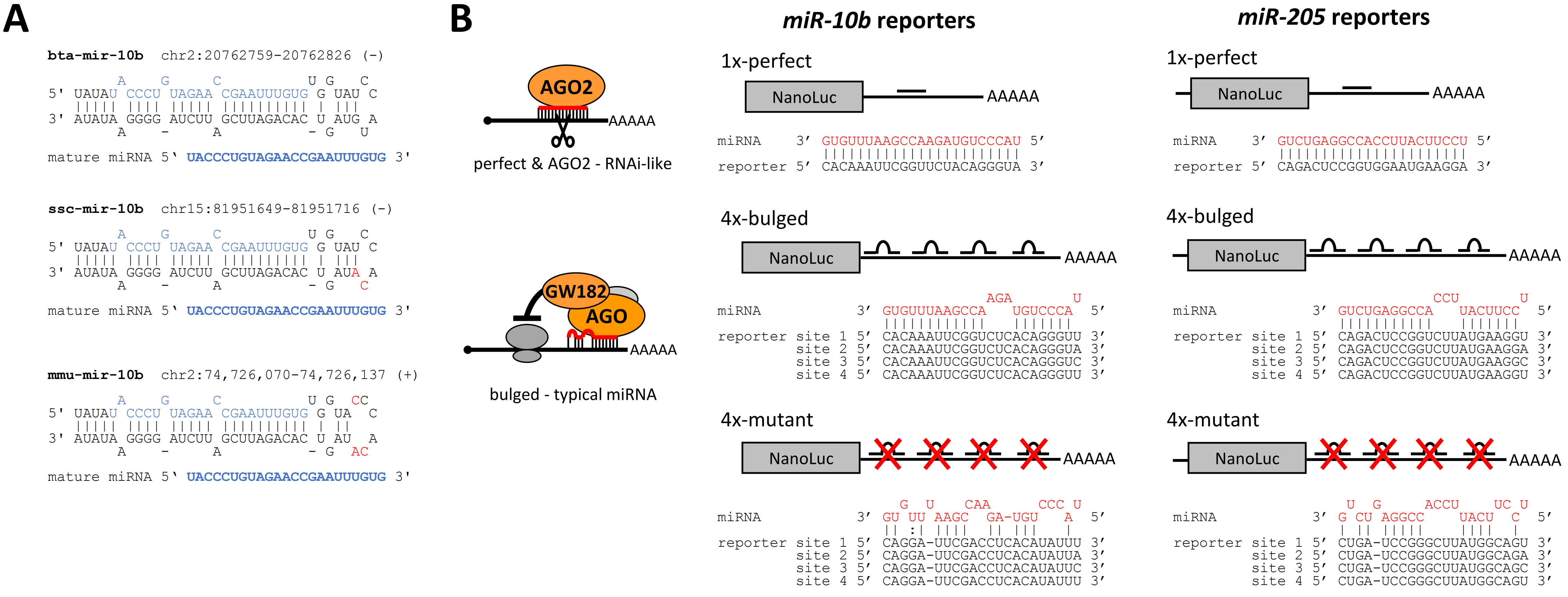
